# Relaxed selection diminishes social memory and expression of host defenses against cuckoos

**DOI:** 10.1101/2024.01.14.575589

**Authors:** Deryk Tolman, Katja Rönkä, Edward Kluen, Juho Jolkkonen, Pietro Di Bari, Renzo Ientile, Daniela Campobello, Rose Thorogood

## Abstract

Behavioral defenses are a key adaptation across animals, but confirming decay under relaxed selection is challenging. If expression involves decision-making, then defenses may only be expressed when the right environmental cues are available (i.e. cryptic plasticity). Even if defenses wane, they could be quickly restored once selection resumes as animals can learn by observing others (i.e. social memory). Here we tested cryptic plasticity and social memory under relaxed selection in a cuckoo host, the Common Reed Warbler, by manipulating perceived parasitism risk with social information in geographically-distinct areas differing in allopatry (100 to 1000 years). Despite predicting social information of a cuckoo vs. a control would elicit otherwise cryptic behavioral defenses, only birds that mobbed model cuckoos prior to manipulation upregulated this defense, and only in the recently allopatric population. Social information also had no effect on low-to-absent rates of foreign egg rejection, a second line of defense common to cuckoo host species. These results contrast with parasitized locations where similar methods provoke mobbing and increase egg rejection ten-fold. Our study therefore suggests that, even when manipulating information to account for reaction norms, behavioral defenses may degrade rapidly under relaxed selection, demonstrating a process facilitating a geographic mosaic of coevolution.

## Introduction

As species’ ranges expand or contract, their exposure to enemies and selection pressures can also change (Gilman et al., 2010). Where selection relaxes (i.e. with allopatry), traits that were previously adaptive are expected to reduce (e.g. due to drift), particularly if maintaining the trait is costly (Lahti et al., 2009). However, while behavioral defenses are common across the animal kingdom (Candolin & Wong, 2015), whether these respond to relaxed selection is rarely tested (Foster, 2013; Lahti et al., 2009). Behaviors are assumed unlikely to be lost due to complex genetic architectures (pleiotropy) and underlying trait variation that can be co-opted in response to other selective pressures (Lahti et al., 2009). However, another key issue is that loss of behavior can be challenging to measure. Behavioral responses depend on prior experience and availability of information about the relative costs or benefits of taking action (i.e. information ecology; Dall et al., 2005). Therefore, the absence of a response does not necessarily indicate a lack of enemy recognition but rather cryptic plasticity (Foster, 2013). Taking an informational approach (Gil et al., 2018; O’Connor et al., 2019) and manipulating cues in allopatry, however, makes it possible to approximate the retention of behavioral reaction norms in the wild.

Social ‘memory’ could play a key role in determining whether behaviors are lost or retained. For example, prey and hosts observe the behavior of others to learn about novel threats (Feeney & Langmore, 2013; Langmore et al., 2012; Rowell et al., 2020) and assess risk of predation or parasitism (Gil et al., 2018), while predators use social information to find suitable prey (Hämäläinen et al., 2022). Therefore, how individuals adjust their behavior in response to social information (i.e. social responsiveness; (de Groot et al., 2023) is likely to affect both their own fitness and the strength of selection on interacting parties (Cantor et al., 2021; Thorogood & Davies, 2012). Social responsiveness may also determine whether populations are resilient to environmental change (Gil et al., 2019; Thorogood et al., 2018), especially as the utility of social information depends on the availability of more-informed ‘demonstrators’. For example, gene flow among populations experiencing variable selection pressures (i.e. trait re-mixing) is thought to determine how quickly a host population responds to shifts in the strength of selection by parasites (Geographic Mosaic Theory of Coevolution; Thompson, 2005). If experienced individuals from ‘hotspots’ of current selection also introduce social information to naïve ‘cold spots’ of relaxed selection, social learning could speed up the recovery of defenses in host populations that are later reinvaded by a previous enemy. However, while recent experimental evolution studies show that social behaviors can diminish very rapidly after selection is removed (Bladon et al., 2023), the role of social cues in maintaining behaviors in ‘cold spots’ is still largely unknown (Lahti et al., 2009; Langmore et al., 2012).

An ideal system to test if behavioral defenses and social responsiveness decline in allopatry, altering the potential for hosts to respond to future (re)invasion, is avian brood parasitism. Virulent brood parasites lay their eggs in the nests of host species and exert strong selection for defensive behaviors such as mobbing or rejecting suspicious eggs. As many brood parasites are aggressive or Batesian mimics (Feeney et al., 2015; Welbergen & Davies, 2011), hosts are also expected to adjust defenses according to local risk to minimize costly mistakes (A. K. Lindholm & Thomas, 2000; Thorogood & Davies, 2013). Furthermore, parasitism can also vary in time and space, with some host populations or species escaping parasitism for many generations (Grim & Stokke, 2016). Over time in allopatry, plasticity is expected to be lost with hosts either becoming resistant to reinvasion (i.e. fixation of defenses) or susceptible to extinction if defenses wane (Soler, 2014) but it is unclear which trajectory will occur as there are examples of both. Some species long separated from brood parasites (i.e. hundreds of thousands to millions of years) show persistent egg rejection when experimentally induced (Bolen et al., 2000; Briskie et al., 1992; Chaumont et al., 2021; Hale & V. Briskie, 2007; Moskát, C. et al., 2002; Rothstein, 2001), however it appears to reduce in others (Briskie et al., 1992; Davies & Brooke, 1989; Soler & Møller, 1990; Yang et al., 2015, 2021). Frontline defenses, such as mobbing of adult brood parasites, also appear to either degrade (Briskie et al., 1992; Hale & V. Briskie, 2007; Lawson et al., 2020, 2023) or be preserved (Briskie et al., 1992; Chaumont et al., 2021; Kuehn et al., 2016) under allopatry. These defenses are not necessarily coupled, for example egg rejection may be expressed while mobbing is not (Hale & V. Briskie, 2007; Thorogood & Davies, 2016). In addition, apparently naïve species can obtain host defenses very rapidly if invaded (Nakamura, 1991) but whether these changes represent the cryptic persistence of defenses in the population, gene flow from parasitized areas, or novel learning is unknown.

Here we use allopatric populations of Common reed warblers (*Acrocephalus scirpaceus*) at the northern- and southernmost extents of their European breeding range to test for the retention of behavioral defenses and social responsiveness to brood-parasitic Common cuckoos (*Cuculus canorus*). Reed warblers colonized Finland ca. 100 years ago (Järvinen & Ulfstrand, 1980), most likely expanding from the southwest (Bergman et al., 2025) and away from regularly parasitized areas in the range core (Avilés et al., 2006). Cuckoos in Finland use different host species, away from reedbeds. In Sicily, reed warblers are likely to have escaped parasitism by cuckoos for much longer – cuckoos only occur on mountainsides while reed warblers are largely restricted to coastal areas, and mentions of both species in historical records are not in association with each other (Arnott, 2007). At parasitized sites in their range-core, reed warblers upregulate mobbing (Campobello & Sealy, 2011; Davies & Welbergen, 2009) and egg rejection (Thorogood & Davies, 2016) defenses after observing their neighbors attack cuckoos. Among populations not currently exposed to parasitism, however, egg rejection rates vary widely (e.g. from 0–40%; (Thorogood & Davies, 2013) and mobbing responses are reduced (e.g. 25% of pairs; (Welbergen & Davies, 2012). However, as in other studies of brood-parasite host defenses in allopatry (see also Rothstein 2001), perceived parasitism risk has not yet been manipulated (e.g. via social information) so the extent to which host defenses reduce under relaxed selection remains unclear.

Therefore, we experimentally manipulated social information and simulated parasitism in field experiments to test if defenses and/or social responsiveness decay in allopatry. We predicted that if defenses are retained, then providing social information should induce plasticity (i.e. social responsiveness) and lead to an increase in mobbing (propensity and intensity) and egg rejection, as previously shown in range-core populations. However, if defenses are not upregulated in response to our manipulated cues, then this would provide evidence that these behaviors are being lost under relaxed selection. Additionally, we assess changes in latency to approach threats at the nest as a more subtle indication of how these hosts perceive cuckoos, since we expect reed warblers to be more attentive when they perceive parasitism risk to be greater (Davies & Welbergen, 2009), and more cautious to approach if they mistake cuckoos for predatory hawks (Thorogood & Davies, 2012; Welbergen & Davies, 2011).

## Materials and Methods

### Study areas and host populations

We studied unparasitised populations of Common reed warblers breeding in May-July around Helsinki, southern Finland (30 discrete reed-lined bays along 100km of coastline, 60.19848N 24.07305E to 60.34039N 25.71162E, in 2019 and 2020) and in the Fiume Ciane nature reserve near Siracusa, Sicily, Italy (37.04285N 15.24273E, in 2022), avoiding re-use of areas across years in Finland to avoid contamination of experience or pseudoreplication. Reed warblers nest at relatively low densities in Helsinki (Tolman et al., 2021) but are more numerous in Fiume Ciane (median distance to closest nest = 36 m, inter-quartile range = 24-53 m). We found nests by cold-searching or monitoring singing males and recorded location and distance to the nearest potential cuckoo vantage point [tree >3 m in height, a possible indirect cue of parasitism risk; (Oien et al., 1996; Welbergen & Davies, 2009)] using handheld GPS (accuracy ± 1 m).

### Experimental protocols

To compare results to previously-studied parasitized populations in the range-core, we adopted published protocols (Davies & Welbergen, 2009; Thorogood & Davies, 2016) as follows. Experiments started on the day the fourth egg was laid (clutch sizes = 3-5 eggs, incubation begins with the penultimate egg; Cramp & Brooks, 1992)). At each nest we measured the change in mobbing response to a cuckoo after presentation of social information of either a cuckoo or control. We used the same 3D-printed model cuckoos (painted by an artist, see Tolman et al., 2021 for details) in Finland and Sicily but controls differed. In Finland we used a commercially-available plastic common magpie (*Pica pica*; Live Decoys) to assess whether changes in mobbing may be due to nest predators or specific to cuckoos, and in Sicily we used plastic female common teal hunting decoys (*Anas crecca;* Flambeau Outdoors), similar to previous range-core experiments using innocuous controls (Davies & Welbergen, 2009). Although this difference precludes comparisons of responses to controls, our focus was on probing the generality of previous findings in the context of relaxed selection, and at parasitized sites there has been similar variation in controls and presentation orders (Campobello & Sealy, 2011; Davies & Welbergen, 2009; Thorogood & Davies, 2012, 2016). To also compare how reed warblers at each site responded to a benign control, we presented teal models at 48 additional nests across the same bays in Finland in 2018-2020, partially overlapping with the timing of the social information experiment. Model cuckoos elicit similar responses as taxidermy specimens (Chen et al., 2022; Welbergen & Davies, 2008) and ours were printed using the same 3D template as independent work (e.g. Marton *et al*. 2019).

To assay mobbing behavior (bill snaps and rasping calls; Welbergen & Davies, 2008), we placed a model adjacent to the nest rim and recorded latency of the first bird to arrive within 3 m after the model was placed and the observer retreated (using a stopwatch, seconds). Latency to arrive was measured to test for changes in nest attentiveness within pairs, which is related to whether birds perceive cuckoos as brood parasites or predatory hawks (Thorogood & Davies, 2012; Welbergen & Davies, 2011). Unrelated variation in latency (e.g. prolonged foraging trips) can be large, but this was minimized since birds were already incubating during experiments and we were mostly interested in changes within pairs but between treatments. We filmed each trial with a small action camera on a pole 1 m from the nest while observing from 3-5 m away, and counted mobbing vocalizations for 5 min (mobbing intensity). Visibility among nests varied, but we only performed experiments when wind conditions were suitable to observe birds moving characteristically through the reeds. Three observers carried out experiments in Finland, each with a different set of models. One observer also performed trials in Sicily. There was high consistency among observers’ counts of bill snaps [ICC: 0.92 (95%CI 0.80–0.98), *p* < 0.001] and rasps [ICC: 0.98 (95%CI 0.95–1.0), *p* < 0.001], as assessed using the Intraclass Correlation Coefficient (ICC) in the *irr* package in R (see Statistical analyses below) from data collected later using 10 videos and blind to field-collected data.

We used simulated neighbor presentations to provide social information 10 m from the focal nest (cf. active neighbors < 40 m in Davies & Welbergen 2009). A previously collected reed warbler nest was attached to reeds at a similar height to the focal nest, with either a model cuckoo or control adjacent to it (treatments assigned alternately). Regardless of the model being presented, we then broadcast one of 5 playback tracks of reed warblers mobbing a model cuckoo [recorded for range-core experiments; (Davies & Welbergen, 2009; Thorogood & Davies, 2016)] for 10 min from a speaker placed under the dummy nest. Focal nests were at least 40 m from other nests or experimental neighbor playbacks to avoid contamination of information. Live mobbing by neighbors does not influence social responsiveness to playbacks (Davies & Welbergen, 2009; Thorogood & Davies, 2012) and simulated vs. active nests previously elicited similar responses to social information treatments in previous experiments (Campobello & Sealy, 2011; Thorogood & Davies, 2016). In Sicily, where densities were higher, we used active neighboring nests for social information presentations in 4 trials (3 cuckoo and 1 teal; < 20m from the focal nest) but none of these focal pairs mobbed a cuckoo. Results were robust as to whether these pairs were included in analyses.

In Finland, we always presented the cuckoo model at the start of the experiment (P1) and after (P3) presentation of social information (SI) of either a magpie control or a cuckoo (Fig. 1A). To avoid pairs in the cuckoo SI treatment being exposed to cuckoos more (i.e. P1, SI, P3) than in the magpie SI treatment (P1 & P3 only), we included a second presentation (P2) of either a cuckoo (magpie SI treatment) or magpie (cuckoo SI treatment) before SI (Fig. 1A). This design also allowed us to compare whether any changes in behavior were due to repeated exposure rather than the SI treatment. To rule out order effects, in Sicily (Fig. 1D) we followed the same design as (Davies & Welbergen, 2009): here each nest was presented with a cuckoo and control (teal) in random order before (P1 & P2) and after (P3 & P4) social information (SI). We did not randomize the order of social information presentations in either study area; however, the likelihood that habituation to models could influence social responsiveness is small as previous experiments exposing reed warblers to additional models before social information was manipulated still detected strong effects (at a parasitized site; Thorogood & Davies 2012). We waited at least 1.5 h between presentations at both sites and conducted trials between 10:00 and 18:00 local time as cuckoos tend to lay in the afternoon. As such, presentations were split over 2 days for each nest.

**Figure 1.**
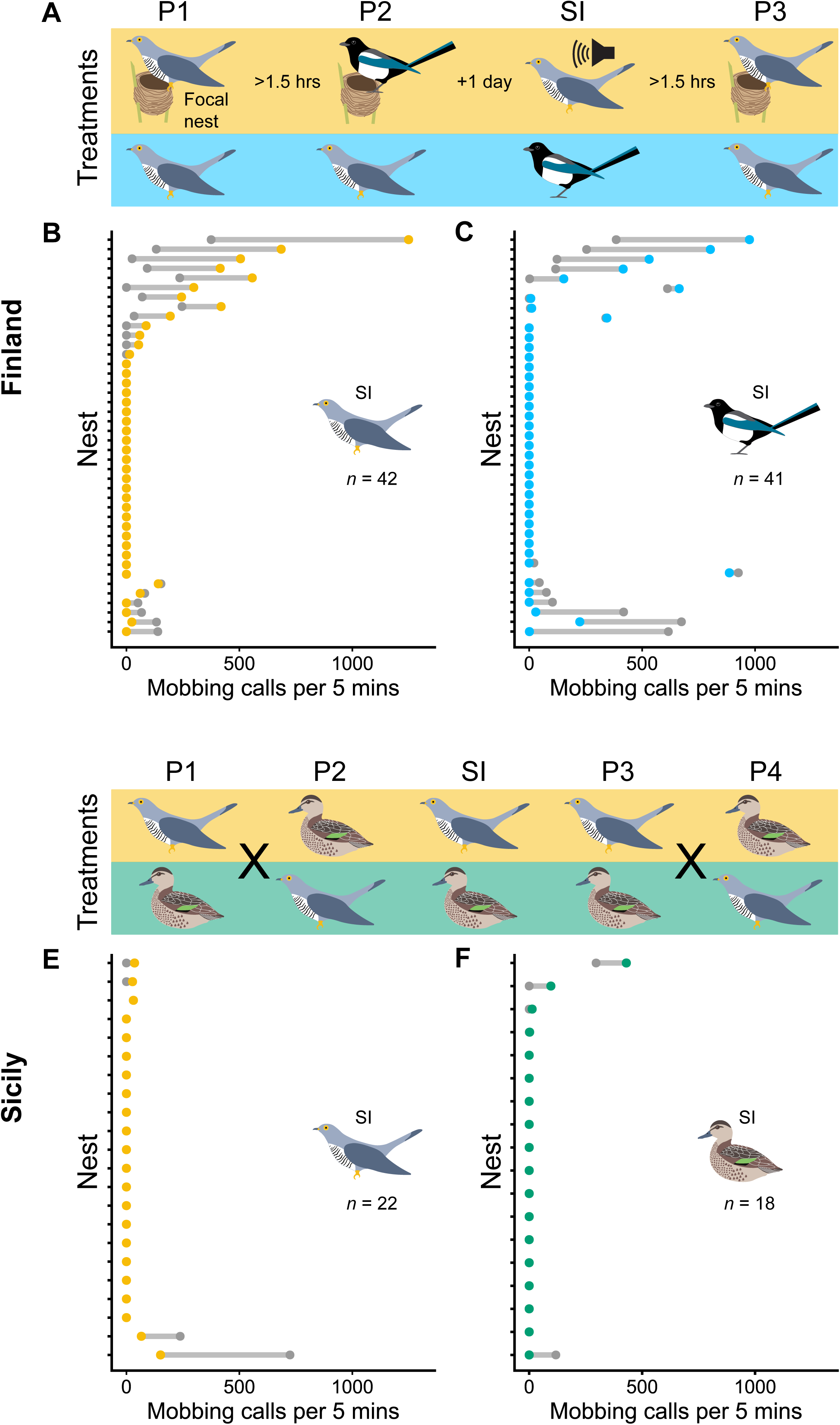
Reed warbler mobbing defences (calls per 5 minutes) in allopatry (Finland: A-C, Sicily: D-F) before (grey dots) and after (coloured dots) experimental manipulation of social information (‘SI’) about cuckoos (B, E) versus a control (C – magpie, F – teal). Details of experimental design are shown in A and D. In Finland (A), the first model presented at nests in Finland (P1) was always a cuckoo, followed by either a cuckoo or magpie (P2) depending on social information treatment (SI), and finished with a cuckoo presentation (P3) for the “after” comparison. In Sicily (D), cuckoo and control model presentation order was alternated before (P1-P2) and after (P3-P4) the social information presentations. **Alt text:** Visual depictions of model presentation order for social information experiments and graphs showing mobbing call rates of reed warblers before and after social information treatments in Finland and Sicily.

In trials where mobbing responses were very weak (i.e. fewer than 20 vocalizations), we later confirmed sounds from video recordings of the trial and excluded five cases where three observers could not discern or agree on sound categorization. We also excluded 4 nests (1 in Finland and 3 in Sicily) where birds failed to appear at one or more cuckoo presentations at their nest (field observations confirmed by video). In total, we analyzed mobbing from 83 Finnish nests (Cuckoo treatment *n =* 42; Magpie treatment *n =* 41) and 40 Sicilian nests (Cuckoo treatment *n =* 22; Teal treatment *n =* 18).

### Measuring egg rejection defenses

After the final model presentation, we colored one egg at random from the focal nest with 40 equally-spaced brown dots using a Staedtler permanent felt tip pen (Lumocolor^®^ 314-7) to measure rejection (as in Thorogood & Davies 2016) of a pseudo reed warbler-gens ‘cuckoo egg’ (Finland: Cuckoo treatment *n =* 24, Magpie treatment *n =* 25; Sicily: Cuckoo treatment *n =* 22, Teal treatment *n =* 18). Due to a low response (see Results), for trials conducted in Finland in 2020 we then used custom 3D-printed eggs with the same dimensions and color (Citadel^©^ blue acrylic paints, custom mixed) as a highly non-mimetic redstart-gens cuckoo egg (Cuckoo treatment *n =* 29, Magpie treatment *n =* 27). In all trials, nests were checked 3-6 days later for removal of the ‘cuckoo egg’ (reed warblers most often reject within 3 days; Davies & Brooke 1988) or evidence of damage (reed warblers were not able to pierce and remove 3D-printed models despite vigorous pecking: 4/25 rejection trials filmed in a subsequent year, unpublished data). Evidence of damage was treated as a rejection attempt. Finally, in Sicily we also assessed non-mimetic egg rejection without any information of cuckoo presence (*n* = 11) and with personal information of parasitism risk (i.e. by presenting a cuckoo model at the nest; *n =* 12) by coloring one egg in the nest entirely with a blue felt-tip pen (Copic Ciao B). Immaculate blue eggs (resembling redstart-gens cuckoo eggs) are highly dissimilar from reed warblers’ and consistently elicit some of the highest rejection rates (often >60%) from this host despite little standardization of color or size across studies (Davies & Brooke, 1988; Dyrcz & Halupka, 2007; Stokke et al., 1999, 2010).

### Statistical analyses

All analyses were conducted in R v.4.3.2 (R Core Team, 2023). First, we used GLMs to check for bias of potential indirect cues (i.e. distance to potential cuckoo vantage points, see Supporting Information S1) and to confirm that initial responses to cuckoos (i.e. P1 in Finland, or P1|P2 in Sicily) were similar in the treatment groups and among years of the study (in Finland). We then used GLMMs with a random effect for nest identity when modelling change in response across multiple presentations (i.e. P1 vs. P3, or P1 vs. P2) as implemented in the package *glmmTMB* (Brooks et al., 2017). To investigate whether social information led to a change in mobbing propensity or intensity, we only included responses to the first and last presentation of a cuckoo at the nest (i.e. P1 and P3 in Finland). However, due to a large proportion of zeroes and lack of suitable fit for a zero-inflated model structure, we adopted a hurdle approach to model mobbing propensity (i.e. binary variable, binomial error structure and logit-link function) and intensity (i.e. number of calls in at least one presentation > 0, quasi-poisson (*nbinom1*) distribution) separately in Finland. For mobbing data from Sicily, we used Fisher’s Exact tests to compare treatment effects on mobbing propensity and quasi-binomial GLMMs to investigate changes in mobbing intensity across all pairs, due to the smaller sample size. To investigate whether social information affected egg rejection decisions, we used a GLM (binomial error structure, logit-link function) which included clutch size as a covariate predictor. Despite data being collected across 30 discrete sites in Finland, including location as a random effect did not improve model fit (Akaike Information Criterion > 2) so it was not part of our final models.

Finally, we investigated latency to approach the threat at the nest during experiments to assess more subtle differences in reaction to cuckoos (i.e. cryptic recognition without a mobbing response). We used separate GLMMs with Tweedie distributions to test if change in latency across presentations differed (i) between treatments and (ii) between mobbing versus non-mobbing pairs as we lacked power to test a Presentation order*SI treatment*Mobbing status 3-way interaction. Nest identity was included as a random effect and pairwise contrasts were estimated using estimated marginal means with Tukey-adjusted p-values. We used QQ-normality plots, residuals vs. predicted plots, and dispersion and outlier tests using package *DHARMa* (Hartig, 2022) to assess assumptions and fits of all models, and we present estimated mean effect sizes (± standard error), *z-*statistics, and *p*-values. Summary tables of all models are available in Supporting Information S2.

## Results

### Mobbing defenses

When first presented with a cuckoo at the nest, 36% (30/83) of pairs in Finland and 13% (5/40) of pairs in Sicily showed a mobbing response (Fig. 1). This initial propensity to mob did not differ between social information (SI) treatment groups (Finland, GLM: estimate = 0.16 ± 0.46, z = −0.35, *p* = 0.73; Sicily, Fisher’s Exact test: *p* = 1.0; Table S2.1 in Supporting Information) or among years of the study in Finland (GLM: estimate = 0.25 ± 0.48, z = 0.52, *p* = 0.60; Table S2.1).

Providing social information about the risk of parasitism in the local area did not induce significantly more reed warblers to mob cuckoos than pairs exposed to information about magpies (in Finland, GLMM Presentation order * SI treatment: estimate = 3.99 ± 2.39, *z* = 1.67, *p* = 0.099, Fig. 1B, Fig. 2A; Table S2.2) or when compared to social information about innocuous teals (in Sicily, across treatments *n* = 10 pairs with calls >0, Fisher’s exact test: *p =* 1.0, Fig. 2A). This lack of effect was not because social information about controls also induced increased mobbing propensity (Finland: Fig. 1C, Fig. 2A; Sicily: Fig. 1F).

**Figure 2.**
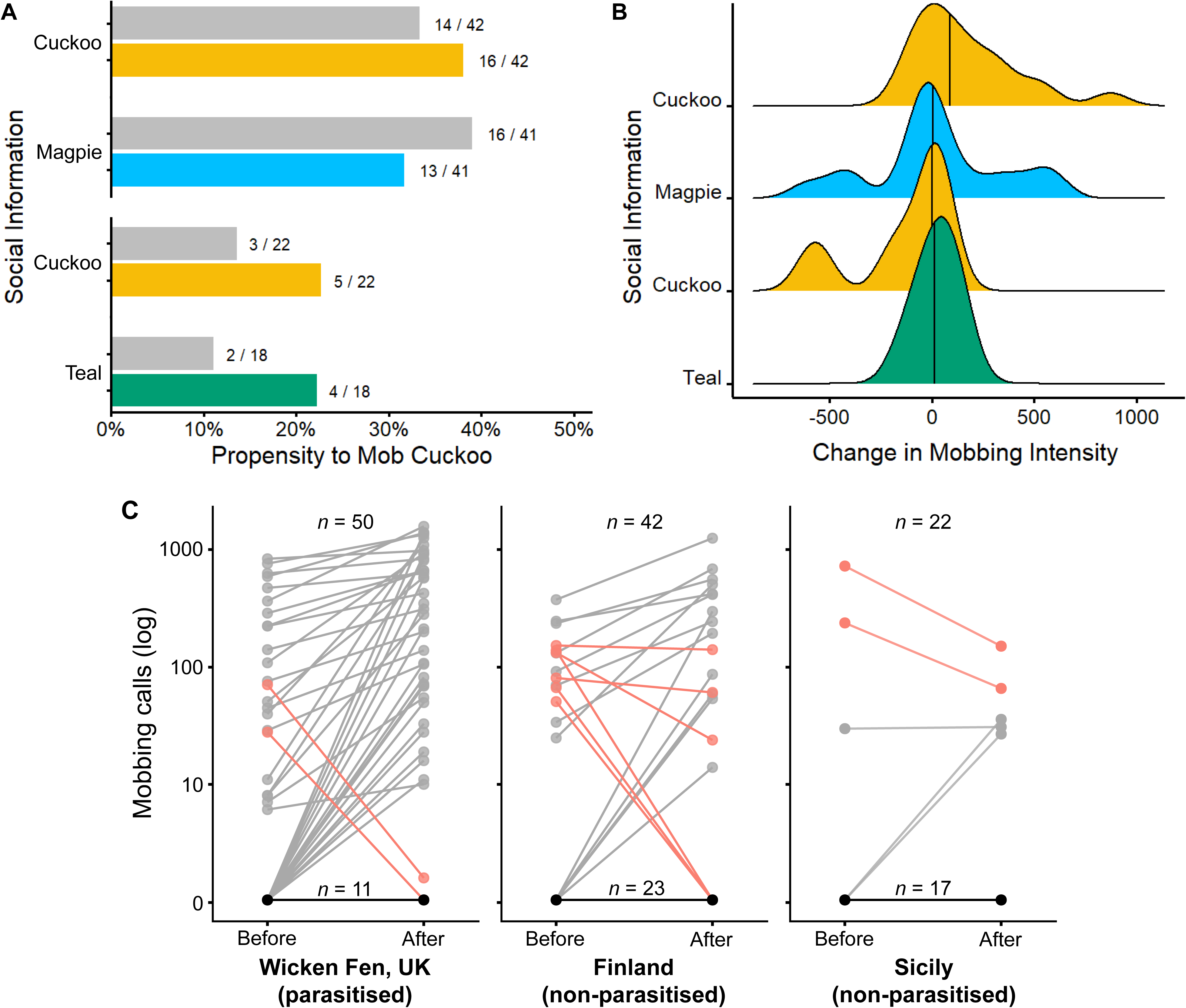
(A) Propensity to mob cuckoos did not change after social information about cuckoos or a control in Finland or Sicily. (B) Mobbing pairs showed social responsiveness (i.e. increased intensity) to cuckoos, but only in Finland (distributions of changes in mobbing intensity, vertical lines = medians). (C) Mobbing reaction norms of reed warblers across parasitism risk (before and after social information of cuckoo activity) were more frequently positive in parasitized Wicken Fen (produced from Davies & Welbergen, 2009 and Thorogood & Davies, 2012 with permission) than in Finland and Sicily. Negative slopes are highlighted in red and sample sizes represent total number of pairs tested (top) and number of slopes that remained at 0 (i.e. did not mob at all; bottom). **Alt text:** Graphs showing mobbing propensity and intensity of reed warblers toward cuckoos before and after social information treatments.

Social information about cuckoos did, however, increase mobbing intensity (i.e. number of signals) but only for pairs who responded in Finland (across treatments *n* = 36 pairs with calls >0, Fig. 1B,C; GLMM Presentation order * SI treatment: estimate = 0.97 ± 0.45, *z* = 2.14, *p* = 0.032, Fig. 2B; Table S2.2). There was no significant change among presentations in the group who received social information about magpies (GLMM estimate = −0.28 ± 0.33, *z* = −0.84, *p* = 0.40, Fig. 2B; Table S2.2). Instead, these pairs were just as likely to increase or reduce mobbing intensity by the third presentation (P3; Fig. 1C). In Sicily, social information had no effect on mobbing intensity (GLMM Presentation order * SI treatment: estimate = −1.20 ± 0.88, *z* = −1.36, *p* = 0.17, Fig. 2B; Fig. 1E, F; Table S2.3). Teal were mobbed at similar intensities between Sicily and Finland (*n* = 96, GLM: estimate = −0.39 ± 0.31, *z* = −1.27, *p* = 0.21; Table S2.5).

### Egg rejection defenses

Reed warblers did not reject ‘mimetic’ brown-spotted eggs in either Finland (Cuckoo treatment = 0/24; Magpie treatment = 0/25; Fig. 3) or Sicily (Cuckoo treatment = 0/21; Teal treatment = 0/19), and there was no rejection of non-mimetic blue eggs in Sicily (no cuckoo presentation = 0/11, after cuckoo presentation at nest = 0/12). Blue eggs were however rejected at low rates in Finland, although this did not differ between social information treatments (GLM: estimate = 0.11 ± 0.64, *z* = 0.17, *p* = 0.97; Fig. 3; Table S2.6).

**Figure 3.**
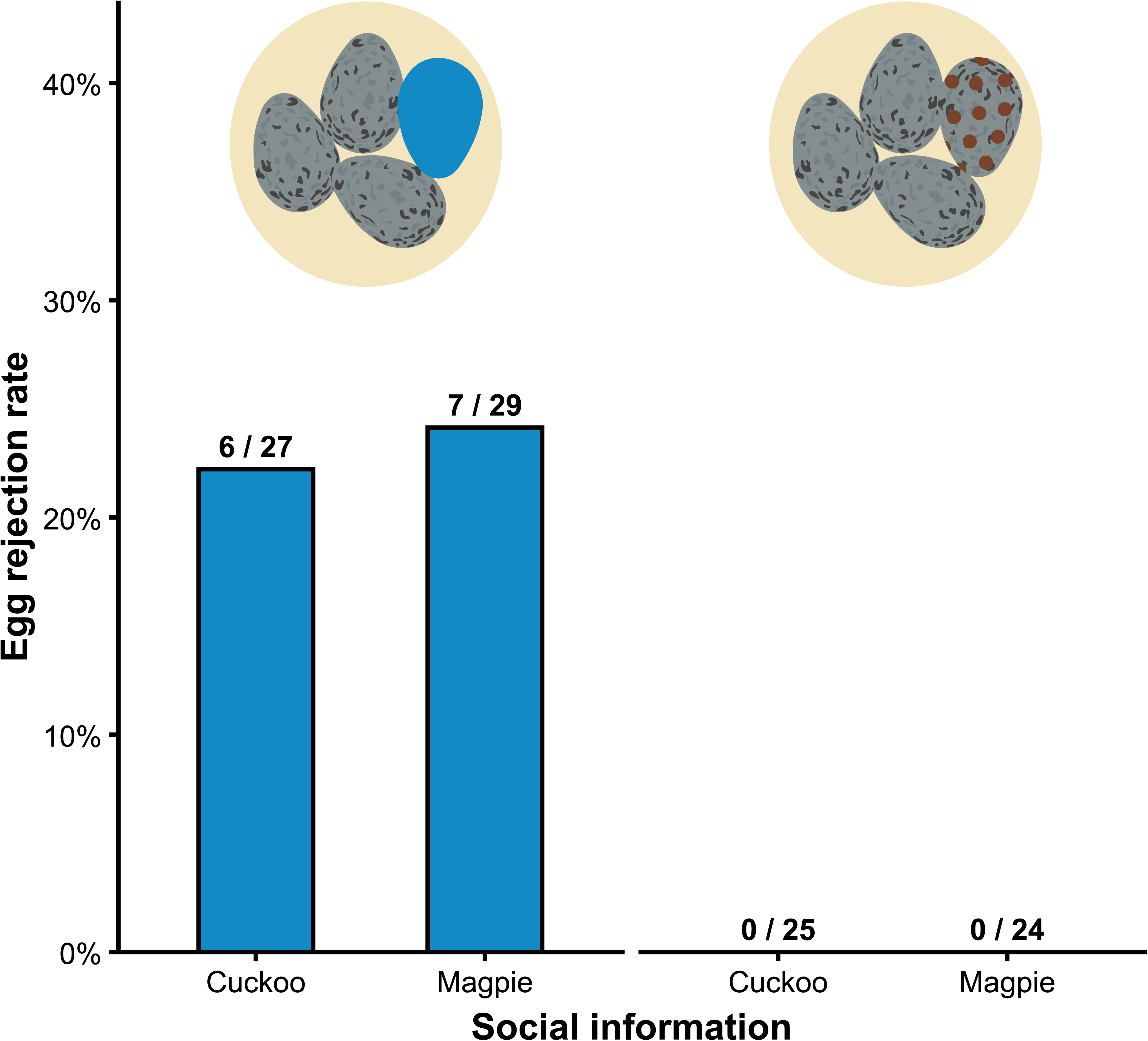
Rejection rates of experimental eggs in Finland (inset illustrations). Blue non-mimetic eggs (left) were rejected at the same rate regardless of whether social information was provided about cuckoos (24%) or magpies (22%). Mimetic brown-spotted eggs (right) were not rejected. **Alt text:** Graph showing egg rejection rates of non-mimetic and mimetic eggs in Finland between social information treatments.

### Recognition of cuckoos vs. response

In Finland, non-mobbing birds were slower initially to approach a cuckoo at their nest than mobbing pairs (P1 data only, GLM: estimate = −0.42 ± 0.19, *z* = −2.29, *p* = 0.022; Fig. 4; Table S2.7) but became faster with subsequent presentations (GLMM Mobbing status * Presentation order, P1 and P3 only, interaction: estimate = 0.54 ± 0.28, *z* = 1.96, *p* = 0.049; Fig. 4; Table S2.8). However, this was not specific to social information because non- mobbers already reduced their latency to approach cuckoos by P2 (P1 vs P2 non-mobbers, GLMM: estimate = −0.44 ± 0.21, z = −2.04, p = 0.10; Table S2.9), and they decreased again after social information of magpies (P2 vs P3 non-mobbers: estimate = −0.43 ± 0.24, z = −1.8, *p* = 0.17; Table S2.9; Fig. 4B). Mobbing pairs showed no change in latency to approach cuckoos across the experiment (P1 vs P3 Cuckoo, GLMM: estimate = 0.10 ± 0.29, z = 0.33, p = 0.94; Table S2.10; Magpie: estimate = 0.17 ± 0.28, z = 0.61, p = 0.81; Table S2.9), and all were more wary of approaching a magpie at the nest than a cuckoo (P2 between treatments, GLM: estimate = 0.72 ± 0.26, *z* = 2.81, *p* = 0.005; Fig. 4A; Table S2.11) even if they never mobbed (mobbing status * threat: estimate = 0.11 ± 0.38, *z* = 0.29, *p* = 0.78; Fig. 4; Table S2.11).

**Figure 4.**
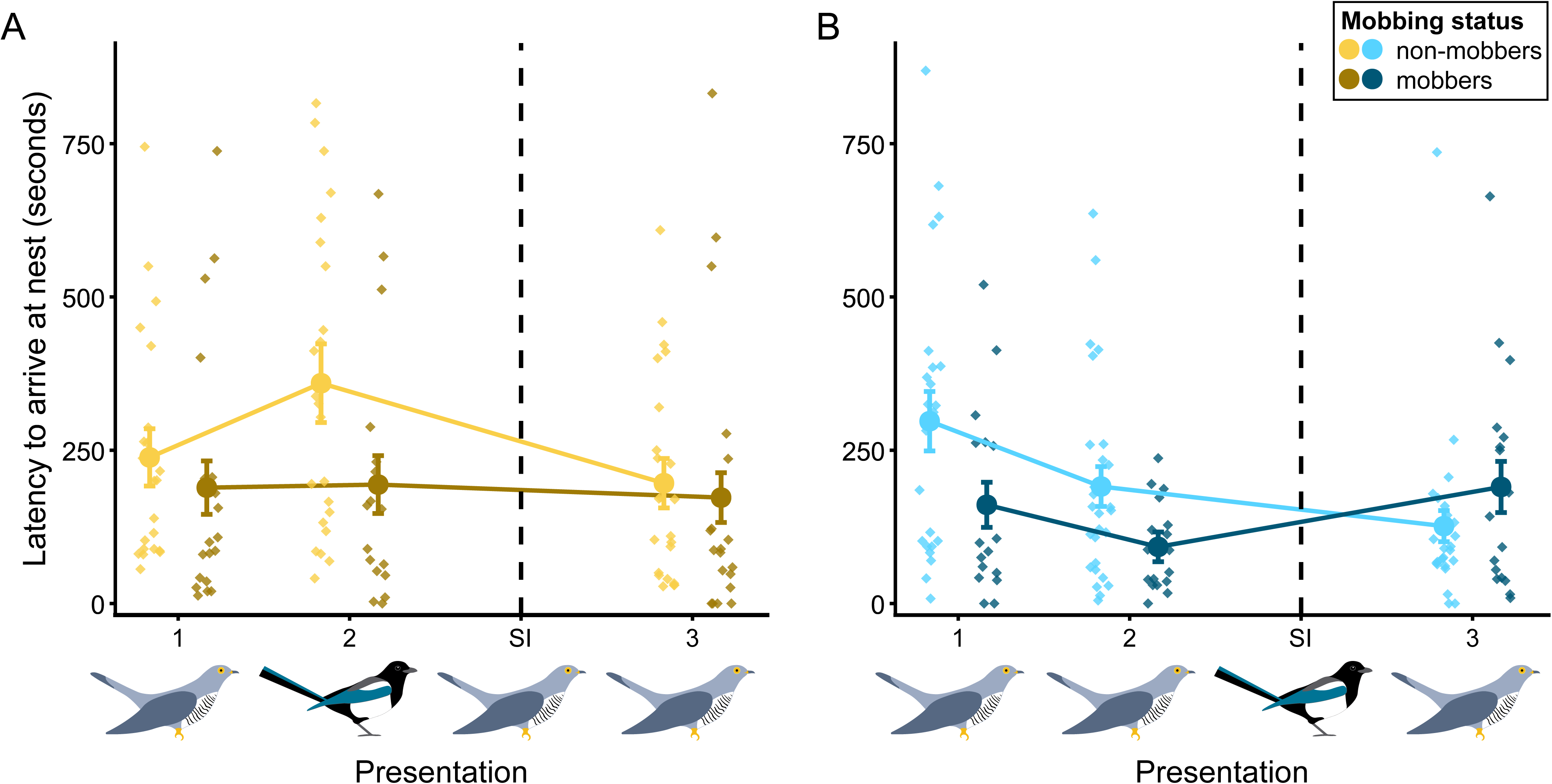
Latency of mobbing and non-mobbing pairs to approach presentations of models at their nest (indicated by icons) before and after social information (SI; dashed line) about (A) cuckoos (yellow) or (B) a magpie control (blue) in Finland. Raw data are plotted as diamonds, with darker symbols indicating pairs that mobbed cuckoos. Predicted mean values (± standard error) from a GLMM are plotted as circles and bars, with connecting lines indicating change across model presentation order (P1-P3). **Alt text:** Graphs showing latencies to approach threat at nest across presentations according to social information treatment and mobbing status.

Reed warblers’ latencies in Sicily did not change over the experiment, either overall (GLMM Presentation order: estimate = 0.20 ± 0.21, *z* = 0.95, *p* = 0.34; Table S2.11) or differently according to social information treatments (GLMM Presentation order * SI treatment: estimate = 0.67 ± 0.42, *z* = 1.58, *p* = 0.12; Table S2.12). They also approached teal as quickly as they did cuckoos (estimate = 0.065 ± 0.168, *z* = 0.39, *p* = 0.67; Table S2.13).

## Discussion

A wealth of prior research in parasitized areas has revealed that reed warblers’ defenses predictably match local brood parasitism risk, including responding to social information of cuckoos by consistently increasing mobbing and egg rejection rates (Campobello & Sealy, 2011; Davies & Welbergen, 2009; Thorogood & Davies, 2016). But, theory also predicts that the loss of defenses in non-parasitized populations is a key part of determining coevolutionary outcomes (Soler, 2014; Thompson, 2005). Although behaviors are assumed hard to lose under relaxed selection, by using social information to simulate increased brood parasitism risk, we found that defenses and social responsiveness of a common host of cuckoos are degrading in areas of allopatry. At the recent range front in Finland, only pairs that mobbed a cuckoo increased their mobbing intensity after receiving social information. The other two-thirds of birds largely remained quiet, although they potentially retained some recognition of cuckoos since both mobbing and non-mobbing pairs approached the nest more quickly than when presented with a magpie, suggesting the decrease in latency of non-mobbers was not purely driven by habituation to disturbance in general, but is specific to the threat. Still, this may reflect avoidance of magpies (Trnka & Požgayová, 2017) rather than awareness of cuckoos and requires further experimental work. Our findings in Finland are enhanced by our results from the Sicilian population, where reed warblers have been in allopatry for longer. There, they showed an even greater reduction in defenses and no evidence of social information use. Importantly, mimetic ‘cuckoo’ eggs remained in nests, and although non-mimetic eggs were rejected at low rates in Finland, social information had no influence on this defense either. This suggests that both allopatric populations would be unlikely to express plasticity or mount significant resistance should cuckoos invade.

Our results are likely evidence for the decay of ancestral defenses rather than a methodological failure to detect responses, for three reasons. First, providing social information experimentally led to an increase in mobbing intensity in Finland, even if not in Sicily. Second, we followed very similar methods to previous studies conducted with reed warblers at parasitized sites, where social information heightened near-absent egg rejection of similarly manipulated eggs ten-fold (Thorogood & Davies, 2016) and induced mobbing of cuckoos (Campobello & Sealy, 2011; Davies & Welbergen, 2009; Thorogood & Davies, 2012), including for pairs that showed no initial response (e.g. 54% of quiet pairs mobbed cuckoos after social information: Davies & Welbergen, 2009). Although there were some differences in our use of controls and presentations of cuckoos at the nest, these represent similar variation to range-core experiments (refs). Third, the differences we detected were specific to cuckoos as responses to a control teal and nest predator (magpie) were similar at our sites and to parasitized areas (Welbergen & Davies, 2008; Campobello & Sealy, 2010; Tolman et al., 2021; Welbergen & Davies, 2012) despite large ecological differences (i.e. climate, nesting density). Some level of cuckoo recognition might be retained in Finland as non-mobbers were slower initially to approach a cuckoo at their nest than mobbing pairs, but became faster with subsequent presentations. If non-mobbers perceived cuckoo models as hawks, we expected that they would instead become more wary (Thorogood & Davies, 2012), but mobbing pairs did not decrease their latency as expected, so it is difficult to conclude that this change in attentiveness is evidence of recognition after all. By using consistent experimental designs to previous studies, testing a large number of reed warblers nesting across a broad range of environmental variation, and considering social responsiveness explicitly, we can therefore be confident that the weak defenses in our allopatric study populations are not the result of cryptic plasticity (Scheiner & Levis, 2021). Instead, they provide compelling evidence that behavioral defenses may sometimes decay in allopatry from cuckoos.

Given that studies of animal culture across a range of taxa demonstrate that the spread of even arbitrary behaviors is possible (Whiten, 2021), it is surprising that we did not detect evidence for social learning among ‘naïve’ reed warblers (i.e. those that did not mob cuckoos). For example, juvenile superb fairy wrens begin to mob cuckoos after observing adults do so (Feeney & Langmore, 2013) and naïve individuals of several species can be trained to recognize novel predators [e.g. introduced species (Rowell et al., 2020)] when paired with mobbing calls. Often social learning of threats requires only one exposure (i.e. similar to our experimental design; reviewed by Crane & Ferrari 2013) and can even be effective when using arbitrary cues (Hämäläinen et al., 2022). Why then did so many birds not mob cuckoos or reject eggs? One possible explanation is that ontogenetic experience or repeated exposure may be necessary before behaviors are expressed, as in the antipredator responses of several fish and mammals (Kelley & Magurran, 2003; Mineka et al., 1984) and personal encounters with avian or piscine brood parasites (Blažek et al., 2018; Hauber et al., 2006; Stokke et al., 2007). Alternatively, but not mutually exclusively, multiple opportunities to learn socially may be needed to elicit a response. While we cannot exclude these possibilities, they seem unlikely to explain the lack of response we detected here: we provided three encounters with cuckoos at or near their nests, yet most birds still did not respond, and under natural conditions naïve pairs are unlikely to repeatedly encounter more responsive neighbors when parasitism rates are low to absent. Perhaps with relaxed selection, there is an initial loss of recognition that then limits hosts’ potential for observational learning.

Temporal and geographical variation in host defenses has typically been explained by plasticity (Langmore et al., 2012; A. Lindholm, 2000; Welbergen & Davies, 2012), particularly in reed warblers where individuals track local parasitism risk and adjust defenses accordingly (Thorogood & Davies, 2013). However, here we found only limited evidence for plasticity in our allopatric populations in response to either indirect environmental or social cues of heightened parasitism risk. This is surprising, given that in a previous study at another unparasitised site of similar length of allopatry to Finland (Llangorse lake, Wales; Lindholm, 1999), reed warblers responded more aggressively to cuckoo models if nesting closer to vantage points for cuckoos (Welbergen & Davies, 2012). We found no evidence for this indirect cue use: instead, birds nesting further from vantage points in Finland were *more* likely to mob, and mobbing in Sicily was unaffected (Supporting Information S1). Nevertheless, of the birds that mobbed, their initial intensities were similar across all three allopatric sites (Finland: 225 ± 43 calls per 5-min, Sicily: 280 ± 120, Llangorse: 281 ± 64; Welbergen & Davies, 2012), suggesting that while the frequency of birds retaining recognition of cuckoos may vary across allopatric populations, reaction norm intercepts have the potential to remain similar. However, comparing reactions to social cues experimentally presented in a similar manner across studies highlights a key difference in allopatry. Previous studies at Wicken Fen, a parasitized site, induced positive social plasticity (*sensu* de Groot et al., 2023) in almost all responding pairs (94.4% across two studies (0+2/18+18 = 5.6%; Davies & Welbergen, 2009, Thorogood & Davies, 2012). At our allopatric sites, on the other hand, reed warblers appeared to be more often ‘confused’, with 30-40% of responding pairs instead reducing mobbing intensity after social information of cuckoos (NEW FIGURE). Interestingly, a similar proportion of negative slopes was observed for birds presented with an unfamiliar cuckoo morph at Wicken Fen (observed only once in 30 years at site; 4/14 pairs declined = 28.6%, Thorogood & Davies 2012). Together this suggests that, as well as a reduction in pairs responding at all, social responsiveness may also weaken in allopatry, potentially leading to the eventual loss of defenses in the absence of cuckoo parasitism (Soler, 2014).

If we were to revisit the Llangorse or Finnish populations in future, would their responses have diminished further and look more like those in Sicily? Quolls’ antipredator behaviors have been found to decay on a predator-free island over even fewer generations than have existed of reed warblers in Finland (Jolly et al., 2018), suggesting gene flow mediates the speed at which behaviors can be lost. There is already some genomic evidence from reed warblers to suggest local adaptation to the strength of parasitism (Smith et al., 2025), but whether the decay in defenses and social responsiveness that we detected is due to genetic change, or a need for additional experience, is yet to be determined. However, it is important to note that the Sicilian population may represent a never-parasitized state where defenses have not previously evolved. The stark lack of defenses in Sicily is therefore an interesting case study of long-term allopatry but, alone, is not evidence for the decay of behaviors in this system. Nevertheless, our findings in Finland are evidence of this, since reed warblers recently arrived in Finland from the parasitized range-core (Bergman et al., 2025). Anti-parasite behaviors are thus assumed to represent the historically selected state, and our cue manipulation should elicit their expression if relevant reaction norms are retained (Campobello & Sealy, 2011; Davies & Welbergen, 2009; Thorogood & Davies, 2012, 2016).

Overall, our results suggest that it may be possible for host populations freed from parasitism to lose expression of defenses, rendering them vulnerable to future reinvasion. The recovery of resistance is unlikely to be enhanced through social transmission, because even if there are individuals who retain recognition of cuckoos, our results show that social responsiveness may also be lost and opportunities for learning would be scarce. Such a reduction of defenses under relaxed selection is one of the key components of geographic mosaic theory explaining the maintenance of coevolution across time and space: rather than the arms race continually escalating, parasites are hypothesized to recurrently invade host populations, or host species, whose defenses have waned in their absence. These so-called coevolutionary cycles (Rothstein, 2001) are thought to only exist during ‘intermediate phases’ of host-parasite interactions, before defenses reach fixation (Soler, 2014), but the rapidity with which a brood parasite host species can acquire (or lose) successful resistance has received little attention. Our results suggest that range expansions and a geographic mosaic of coevolution may moderate the arms race, with defenses decaying within even relatively short time frames (e.g. approximately 35 generations in Finland; Ceresa et al., 2015), allowing the longer-term persistence of parasites until host switching becomes more favorable (Nuismer et al., 2003). As both social information use and behavioral defenses against enemies are common across the animal kingdom, and species’ ranges are changing rapidly, it is now crucial to determine how socially-mediated plasticity degrades under relaxed selection in the wild.

## Author Contributions

**Deryk Tolman:** conceptualization, data curation, formal analysis, investigation, methodology, project administration, supervision, validation, visualization, writing - original draft, writing - review & editing; **Katja Rönkä:** conceptualization, data curation, investigation, methodology, project administration, resources, writing - review & editing; **Edward Kluen:** conceptualization, data curation, investigation, methodology, project administration, writing - review & editing; **Juho Jolkkonen:** investigation, methodology, writing - review & editing; **Pietro Di Bari:** investigation, methodology, writing - review & editing; **Renzo Ientile:** investigation, project administration, resources, writing - review & editing; **Daniela Campobello:** investigation, project administration, resources, supervision, writing - review & editing; **Rose Thorogood:** conceptualization, data curation, formal analysis, funding acquisition, investigation, methodology, project administration, resources, supervision, validation, visualization, writing - review & editing.

## Acknowledgements

We thank field assistants Julius Mäkinen, Anna Tuominen, Anna Välkki, Purabi Deshpande, Linnea Kivelä, and Nora Bergman for their essential help with nest searching and monitoring, Lukas Wolfram for confirming responses to model eggs from videos, Tuomas Pärnänen for designing and printing eggs, Steve Collett for painting model cuckoos, Merve Ünlü for bird illustrations, and Gloria Murari for figure aesthetics.

We are grateful to private landowners, Uusimaa municipalities, and Metsähallitus (permission no. MH1100/2018/06.06.02) for allowing access to reed beds in Finland, and Giuseppe Mammino and staff of the Fiume Ciane Nature Reserve (Libero Consorzio Comunale di Siracusa) for facilitating our work in Sicily. We also thank the editors and two anonymous reviewers for helpful comments which improved this manuscript.

