## Supplementary Information for "Relaxed selection diminishes social memory and expression of host defenses against cuckoos"

**This PDF file includes:**

Supporting text S1
Figure S1
References for S1
Tables S2.1 to S2.13

**Supporting Information S1. Ruling out the use of indirect cues of parasitism risk.**

Methods

Several previous studies have found reed warblers' mobbing defences to be correlated to distance to the nearest potential perch for cuckoos to observe the location and behaviour of potential hosts (e.g. Oien *et al.* 1996), including at another allopatric site (Welbergen & Davies 2012). We used the same criteria and quantified this as distance to the nearest tree greater than 3 m in height, measured using handheld GPS (accuracy ± 1 m). Prior to other analyses, we confirmed there was no bias between the two treatment groups in the distribution of distance to potential cuckoo perches (gamma GLMs with treatment group, Finland: estimate = -0.001 ± 0.003, z = -0.41, p = 0.69; Sicily: estimate = -0.001 ± 0.007, z = -0.14, p = 0.89). We then tested whether this cue of parasitism risk was used by birds in our study using GLMs with binomial error structures.

Results

We found birds in Finland to be more likely to mob cuckoos the further they were from these potential cuckoo vantage points (estimate = 0.023 ± 0.010, $z$ = 2.31, $p$ = 0.021; Fig. S1A), suggesting that this was not used as an indirect cue of parasitism risk but that there could be some other ecological variable involved in explaining variation in mobbing. There was no trend in either direction for birds in Sicily (0.0007 ± 0.0155, $z$ = 0.047, $p$ = 0.96; Fig. S1B).

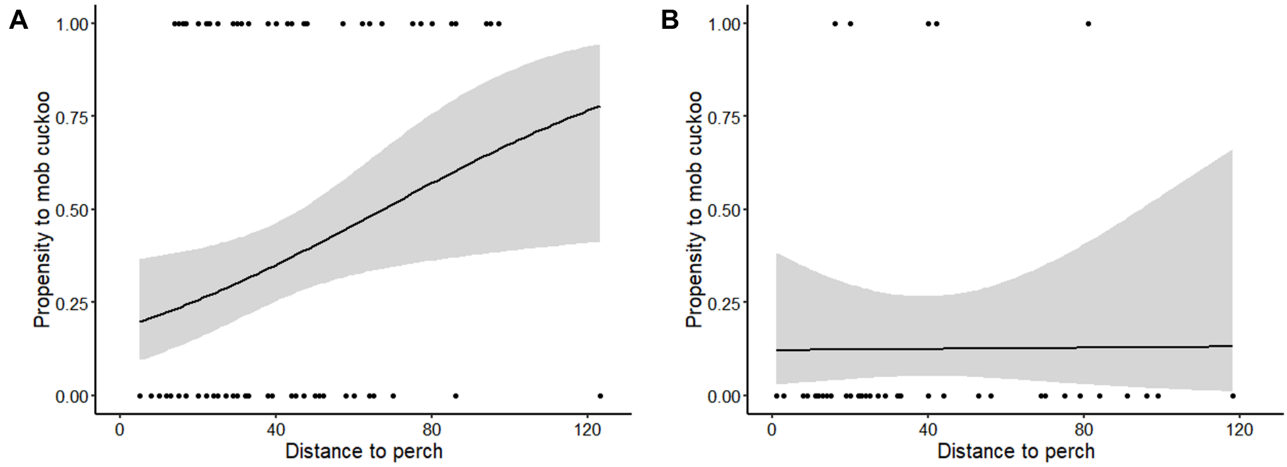


**Figure S1.** Relationship between distance to potential cuckoo perch and propensity to mob a cuckoo model at P1 (i.e. before social information treatments applied) in (A) Finland and (B) Sicily. Raw data and binomial trend lines (± SE) are plotted.

**Supporting Information S2. Summary tables of statistical models.**

**Table S2.1.** Mobbing propensity at P1 between treatments and years in Finland

| | **Mobbing propensity** | | | |
|---|---|---|---|---|
| *Predictors* | *Log-Odds* | *SE* | *z* | *p* |
| (Intercept) | -0.52 | 0.37 | -1.40 | 0.160 |
| SI treatment | -0.24 | 0.46 | -0.52 | 0.606 |
| Year | 0.19 | 0.48 | 0.39 | 0.694 |

**Table S2.2.** Mobbing propensity and intensity towards cuckoos in Finland between presentations and social information treatments.

| | **Mobbing propensity** | | | | **Mobbing intensity** | | | |
|---|---|---|---|---|---|---|---|---|
| *Predictors* | *Log-Odds* | *SE* | *z* | *p* | *Log-Mean* | *SE* | *z* | *p* |
| (Intercept) | -7.08 | 1.47 | -4.83 | **<0.001** | 5.35 | 0.34 | 15.63 | **<0.001** |
| Presentation order | -2.01 | 1.43 | -1.40 | 0.161 | -0.28 | 0.33 | -0.84 | 0.399 |
| SI treatment | -1.12 | 1.66 | -0.68 | 0.499 | -0.80 | 0.43 | -1.85 | 0.064 |
| Presentation order * SI treatment | 3.01 | 1.83 | 1.65 | 0.099 | 0.97 | 0.45 | 2.14 | **0.032** |
| **Random Effects** | | | | | | | | |
| $\sigma^2$ | 3.29 | | | | 0.01 | | | |
| $\tau_{00}$ | 184.74 $_{NestID}$ | | | | 0.70 $_{NestID}$ | | | |
| N | 83 $_{NestID}$ | | | | 36 $_{NestID}$ | | | |

**Table S2.3.** Mobbing intensity towards cuckoos in Sicily between presentations and social information treatments.

| | Mobbing intensity | | | |
|---|---|---|---|---|
| *Predictors* | *Log-Mean* | *SE* | *z* | *p* |
| (Intercept) | -1.88 | 2.17 | -0.86 | 0.388 |
| Presentation order | -0.50 | 0.63 | -0.80 | 0.422 |
| SI treatment | -0.48 | 1.57 | -0.31 | 0.759 |
| Presentation order * SI treatment | 1.20 | 0.88 | 1.36 | 0.173 |
| **Random Effects** | | | | |
| $\sigma^2$ | 0.01 | | | |
| $\tau_{00\ NestID}$ | 22.73 | | | |
| $N_{NestID}$ | 40 | | | |

**Table S2.4.** Mobbing propensity and intensity towards cuckoos between Sicily and Finland.

| | Mobbing propensity | | | | Mobbing intensity | | | |
|---|---|---|---|---|---|---|---|---|
| *Predictors* | *Log-Odds* | *SE* | *z* | *p* | *Log-Mean* | *SE* | *z* | *p* |
| (Intercept) | -0.19 | 0.22 | -0.87 | 0.383 | 5.57 | 0.33 | 17.05 | **<0.001** |
| Location | -1.03 | 0.42 | -2.45 | **0.014** | -1.44 | 0.72 | -2.01 | **0.044** |

**Table S2.5.** Mobbing intensity towards teal between Sicily and Finland.

| | Mobbing intensity | | | |
|---|---|---|---|---|
| *Predictors* | *Log-Mean* | *SE* | *z* | *p* |
| (Intercept) | 4.22 | 0.33 | 12.47 | **<0.001** |
| Location | -0.34 | 0.31 | -1.1 | 0.283 |

**Table S2.6.** Rejection of blue eggs according to social information treatment and final clutch sizes.

| | **Egg rejection** | | | |
|---|---|---|---|---|
| *Predictors* | *Log-Odds* | *SE* | *z* | *p* |
| (Intercept) | -7.55 | 2.83 | -2.67 | **0.008** |
| SI treatment | 0.02 | 0.67 | 0.04 | 0.972 |
| Clutch size | 1.47 | 0.63 | 2.31 | **0.021** |

**Table S2.7.** Approach time at P1 and P3 according to mobbing status (did, or did not mob, a cuckoo at either, or both, P1 or P3) in Finland.

| | **Approach time P1** | | | | **Approach time P3** | | | |
|---|---|---|---|---|---|---|---|---|
| *Predictors* | *Estimates* | *SE* | *z* | *p* | *Estimates* | *SE* | *z* | *p* |
| (Intercept) | 5.59 | 0.12 | 48.32 | **<0.001** | 5.08 | 0.14 | 36.61 | **<0.001** |
| Mobbing status | -0.42 | 0.18 | -2.29 | **0.022** | 0.12 | 0.21 | 0.58 | 0.562 |

**Table S2.8.** Approach time according to presentation type and mobbing status in Finland.

| | **Approach time** | | | |
|---|---|---|---|---|
| *Predictors* | *Estimates* | *SE* | *z* | *p* |
| (Intercept) | 5.59 | 0.12 | 46.99 | **<0.001** |
| Mobbing status | -0.42 | 0.19 | -2.21 | **0.027** |
| Presentation order | -0.51 | 0.18 | -2.87 | **0.004** |
| Mobbing status * Presentation order | 0.54 | 0.28 | 1.96 | **0.049** |

**Table S2.9.** Approach time to threat at nest according to mobbing behaviour (mobber vs. non-mobber) and presentation order (P1, P2, P3) for magpie social information treatment in Finland.

| | Approach time | | | |
|---|---|---|---|---|
| *Predictors* | *Estimates* | *SE* | *z* | *p* |
| (Intercept) | 5.44 | 0.18 | 29.44 | <0.001 |
| Mobber | -0.24 | 0.28 | -0.85 | 0.398 |
| Presentation P2 | 0.41 | 0.23 | 1.75 | 0.080 |
| Presentation After | -0.19 | 0.25 | -0.76 | 0.448 |
| Mobber * Presentation P2 | -0.38 | 0.38 | -1.01 | 0.313 |
| Mobber * Presentation After | 0.09 | 0.38 | 0.24 | 0.808 |
| **Random Effects** | | | | |
| σ2 | 7.87 | | | |
| τ00 NestID | 0.07 | | | |
| N NestID | 42 | | | |

**Pairwise contrasts**

**Presentation contrasts within non-mobbers**

| *Contrast* | *Estimates* | *SE* | *z* | *p* |
|---|---|---|---|---|
| Before – P2 | -0.41 | 0.23 | -1.75 | 0.185 |
| Before – After | 0.19 | 0.25 | 0.76 | 0.729 |
| P2 – After | 0.60 | 0.24 | 2.50 | 0.034 |

**Presentation contrasts within mobbers**

| *Contrast* | *Estimates* | *SE* | *z* | *p* |
|---|---|---|---|---|
| Before – P2 | -0.03 | 0.29 | -0.11 | 0.993 |
| Before – After | 0.10 | 0.29 | 0.33 | 0.941 |
| P2 – After | 0.13 | 0.30 | 0.43 | 0.903 |

**Mobber contrasts within presentation**

| *Contrast* | *Estimates* | *SE* | *z* | *p* |
|---|---|---|---|---|
| Mobber – Non-mobber (Before) | -0.24 | 0.28 | -0.85 | 0.398 |
| Mobber – Non-mobber (P2) | -0.62 | 0.28 | -2.21 | 0.027 |
| Mobber – Non-mobber (After) | -0.14 | 0.29 | -0.49 | 0.622 |

**Table S2.10.** Approach time to threat at nest according to mobbing behaviour (mobber vs. non-mobber) and presentation order (P1, P2, P3) for cuckoo social information treatment in Finland.

| | Approach time | | | |
|---|---|---|---|---|
| *Predictors* | *Estimates* | *SE* | *z* | *p* |
| (Intercept) | 5.67 | 0.16 | 36.13 | <0.001 |
| Mobber | -0.62 | 0.26 | -2.38 | 0.017 |
| Presentation P2 | -0.44 | 0.21 | -2.04 | 0.041 |
| Presentation After | -0.87 | 0.23 | -3.73 | <0.001 |
| Mobber * Presentation P2 | -0.10 | 0.37 | -0.27 | 0.787 |
| Mobber * Presentation After | 1.04 | 0.36 | 2.86 | 0.004 |
| **Random Effects** | | | | |
| σ2 | 7.30 | | | |
| $\tau 00_{NestID}$ | 0.05 | | | |
| $N_{NestID}$ | 45 | | | |

**Pairwise contrasts**

**Presentation contrasts within non-mobbers**

| *Contrast* | *Estimates* | *SE* | *z* | *p* |
|---|---|---|---|---|
| Before – P2 | 0.44 | 0.21 | 2.04 | 0.102 |
| Before – After | 0.87 | 0.23 | 3.74 | <0.001 |
| P2 – After | 0.43 | 0.24 | 1.81 | 0.166 |

**Presentation contrasts within mobbers**

| *Contrast* | *Estimates* | *SE* | *z* | *p* |
|---|---|---|---|---|
| Before – P2 | 0.54 | 0.31 | 1.76 | 0.183 |
| Before – After | -0.17 | 0.28 | -0.61 | 0.813 |
| P2 – After | -0.71 | 0.30 | -2.36 | 0.048 |

**Mobber contrasts within presentation**

| *Contrast* | *Estimates* | *SE* | *z* | *p* |
|---|---|---|---|---|
| Mobber – Non-mobber (Before) | -0.62 | 0.26 | -2.38 | 0.017 |
| Mobber – Non-mobber (P2) | -0.72 | 0.28 | -2.53 | 0.012 |
| Mobber – Non-mobber (After) | 0.43 | 0.27 | 1.55 | 0.120 |

**Table S2.11.** Approach time at P2 between social information treatments (cuckoo model vs magpie model) in Finland.

| | Approach time | | | |
|---|---|---|---|---|
| *Predictors* | *Estimates* | *SE* | *z* | *p* |
| (Intercept) | 5.25 | 0.16 | 32.67 | **<0.001** |
| Threat | 0.63 | 0.22 | 2.84 | **0.005** |
| Mobbing status | -0.73 | 0.28 | -2.56 | **0.010** |
| Threat * Mobbing status | 0.11 | 0.38 | 0.29 | 0.776 |

**Table S2.12.** Approach time to a cuckoo according to presentation order (before vs. after social information) and social information treatment group (cuckoo vs. control) in Sicily.

| | **Approach time (with interaction)** | | | | **Approach time (without interaction)** | | | |
|---|---|---|---|---|---|---|---|---|
| *Predictors* | *Estimates* | *SE* | *Z* | *p* | *Estimates* | *SE* | *z* | *p* |
| (Intercept) | 4.68 | 0.25 | 18.60 | **<0.001** | 4.55 | 0.24 | 19.17 | **<0.001** |
| Presentation order | -0.07 | 0.27 | -0.26 | 0.793 | 0.20 | 0.21 | 0.95 | 0.343 |
| SI treatment | -0.74 | 0.39 | -1.90 | 0.058 | -0.39 | 0.32 | -1.20 | 0.229 |
| Presentation order * SI treatment | 0.67 | 0.42 | 1.58 | 0.115 | | | | |
| **Random Effects** | | | | | | | | |
| $\sigma^2$ | 0.55 | | | | 0.57 | | | |
| $\tau_{00}$ | 0.55 $_{NestID}$ | | | | 0.54 $_{NestID}$ | | | |
| N | 40 $_{NestID}$ | | | | 40 $_{NestID}$ | | | |

**Table S2.13.** Approach time to the nest depending on threat type (cuckoo vs. innocuous teal) in Sicily.

| | **Approach time** | | | |
|---|---|---|---|---|
| *Predictors* | *Estimates* | *SE* | *z* | *p* |
| (Intercept) | 4.21 | 0.20 | 21.06 | **<0.001** |
| Threat | 0.07 | 0.17 | 0.39 | 0.698 |
| **Random Effects** | | | | |
| $\sigma^2$ | 0.36 | | | |
| $\tau_{00}$ $_{NestID}$ | 0.90 | | | |
| N $_{NestID}$ | 40 | | | |
